# Biological Embedding of Social Experience Over Genomic and Neural Time Scales

**DOI:** 10.64898/2026.11.17.767304

**Authors:** Tianfei Peng, Amy C. Cash Ahmed, Melissa E. Flores, Naroa Legarra-Marcos, Mikel Hernaez, Autumn K. McLane-Svoboda, Summer B. McLane-Svoboda, Debajit Saha, Gene E. Robinson

## Abstract

The “Neurons for Today, Genes for Tomorrow” model (NTGT) predicts how the brain’s neural and molecular control systems interact to regulate biological embedding, whereby social experience alters the brain to shape future behavior. We report here on the first test of this model, using honey bee queen retinue pheromone (QRP) and behavioral, single-nucleus transcriptomic, and electrophysiological analyses. Consistent with theory, weaker exposure triggered a simple attraction response and transcriptomic changes primarily in the antennal lobes (AL), whereas stronger exposure induced transcriptomic changes in the mushroom bodies, a brain region affecting more complex behavior. In addition, AL electrophysiological responses to QRP were enhanced after stronger exposure, also consistent with theory. We also showed how individual differences in biological embedding can be explained by NTGT, with individual differences in brain gene regulatory network topology, especially in specific sensory cell clusters, supporting a new mechanistic framework for behavioral regulation.

## Introduction

Biological embedding, whereby social experience alters the brain to shape future behavior, occurs across diverse animal lineages and has important implications for human mental health^1,2^. Social experience can differ in valence, ranging from affiliative cues that promote approach or cooperation to agonistic cues associated with threat, competition, or avoidance. These different forms of social experience have distinct consequences for behavior, physiology, and ultimately fitness^3^. The perception of negatively valenced social stimuli is also highly relevant to human psychiatric disorders associated with trauma, as even brief environmental encounters can produce lasting effects^4^. The role of neural signaling in biological embedding has been studied extensively^5,6^, but much less is known about the role played by brain transcriptomic changes.

Although brain function has traditionally been understood through the specialized properties of neurons and the circuits they form, recent transcriptomic studies suggest that behavior is also shaped by dynamic changes in brain gene expression and rewiring of brain gene regulatory networks (GRNs) that link neural activity and experience to changes in behavior^7–9^. Across diverse animal species, behavioral states are associated with coordinated changes in thousands of brain-expressed genes, and these expression patterns can sometimes predict behavior^10–13^. But often these transcriptomic changes are detected after a change in behavioral state is observed^14,15^, raising the question of functional significance.

The verbal model, “Neurons for Today, Genes for Tomorrow” (NTGT), was developed to address this puzzle, providing a framework for understanding how neural and molecular layers of regulation operate across different timescales^16,17^. NTGT starts with the well-established understanding that social experience first engages sensory, then higher-order, neural circuits that trigger behavioral change, often in concert with activity-dependent changes in brain gene expression. NTGT predicts that weaker stimuli (reflecting more fleeting experience) trigger smaller-scale molecular changes and do not necessarily change future behavior, while stronger stimuli trigger larger molecular changes that are more likely to shape future behavior.

NTGT seeks to explain links between immediate neural responses and molecular processes with longer latency by which social interactions become biologically embedded. This is similar to memory temporal dynamics, with short-term memory involving changes in existing synapses and long-term memory requiring new gene expression and protein synthesis^18^. This is another reflection of the commonalities between instinctive and learned behaviors at the neural and molecular levels^19^.

The goal of this study was to test NTGT predictions in order to gain a better understanding of how neural and molecular control systems work together to regulate behavior. We used the Western honey bee and its diverse responses to queen retinue pheromone (QRP) as our test system.

QRP is a complex blend of nine components produced by several exocrine glands in the queen that together act as both a releaser and a primer pheromone. QRP affects a range of physiological and behavioral processes in worker bees, the members of the bee society responsible for all aspects of colony function aside from reproduction. As a releaser pheromone, QRP elicits immediate behavioral responses such as promoting retinue behavior, with worker bees surrounding the queen to feed and groom her^20^. As a primer pheromone, QRP induces longer-term physiological changes that ultimately alter behavior, such as raising the behavioral response threshold to sucrose, which is related both to rates of behavioral maturation and foraging decisions in the field^21^.

QRP offers an experimentally tractable system in which stimulus exposure can be precisely controlled, and we used different QRP exposure durations to represent different strengths of social stimulus. The releaser and primer effects of QRP allowed us to examine social regulation across different timescales, providing an ideal system to test the predictions of NTGT.

## Results

### Experiment 1: QRP effects on behavior vary with the strength of social stimulus

We first tested the general prediction of NTGT that the strength of social stimuli can shape future behavior. We used adult worker bees derived from queens that were each instrumentally inseminated with a single drone (SDI) to minimize genetic variation across treatment groups and compared age-matched groups exposed to different strengths of QRP stimulus. Individuals in the short-exposure group (SE group) were kept without QRP from adult emergence until 3 days of age, followed by a 1-hour QRP exposure representing a weaker social stimulus. Individuals in the long-exposure group (LE group) were exposed continuously to QRP from emergence until 3 days of age, representing a stronger social stimulus. We then compared the behavioral responses of the SE and LE groups with those of control individuals that were never exposed to QRP (nE group).

QRP had a significant effect on the retinue response (N = 120 individuals per group from each colony, 4 colonies), but the effect did not depend on QRP stimulus strength (Figure 1A). QRP treatment significantly increased the number of individuals showing retinue behavior regardless of exposure strength: individuals in both SE and LE groups showed higher responses relative to the nE group (LE vs nE: z = 5.68, p < 0.0001; SE vs nE: z = 5.25, p < 0.0001), while responses did not differ between the SE and LE groups (z = 0.91, p = 0.63).

**Figure 1.**
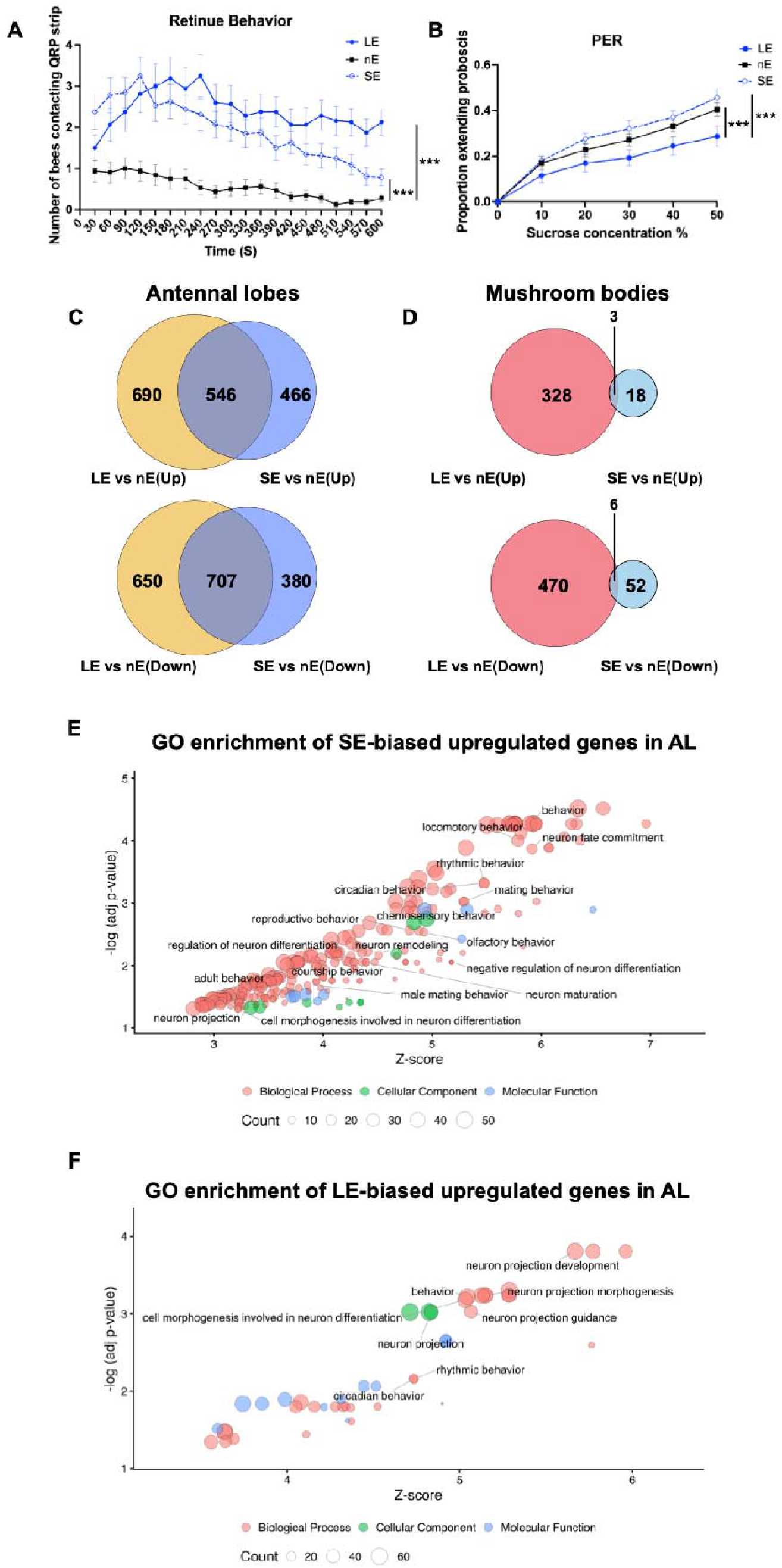
Behavioral and transcriptomic responses to different strengths of Queen Retinue Pheromone stimulus. **(A)** Retinue behavior of bees exposed to a shorter QRP stimulus (1h, SE) or a longer QRP stimulus (3 days, LE) or no exposure (control, nE). The number of bees contacting the QRP strip was recorded every 30 s during a 10-min assay. Data were collected in 2024 from 4 colonies, with 480 bees tested per treatment group. SE and LE individuals showed stronger retinue responses than nE individuals (***p < 0.001 for each comparison). Individuals in these experiments were derived from queens each instrumentally inseminated with semen from a single drone; similar results were obtained for individuals derived from naturally mated queens (Figure S1A). QRP stimulus strength was varied by controlling length of exposure; a stronger stimulus was the result of a longer exposure. **(B)** Proboscis extension response (PER) to increasing sucrose concentrations (0%–50%) in groups of SE, LE, and nE individuals, shown as the proportion of bees extending the proboscis. N = 383 (SE), 383 (LE), and 382 (nE) bees from 4 colonies (2024). LE individuals were less responsive to sucrose than SE and nE individuals (***p < 0.001 for each comparison). Individuals in these experiments were derived from queens each instrumentally inseminated with semen from a single drone; similar results were obtained for individuals derived from naturally mated queens (Figure S1B). QRP stimulus strength was varied as in Figure 1A. **(C and D)** Venn diagrams showing overlap between differentially expressed genes (DEGs) following weaker (1h, SE) or stronger (3 days, LE) QRP stimulus exposure relative to untreated controls (nE) in the antennal lobes (AL; C) and mushroom bodies (MB; D). Upregulated and downregulated genes are shown separately. Numbers indicate the numbers of genes unique to each exposure duration or shared between the two comparisons. QRP stimulus strength was varied as in Figure 1A. **(E and F)** GO enrichment analysis of the AL upregulated genes unique to SE (E) or LE (F). Each point represents an enriched GO term. The x-axis shows enrichment Z-score, and the y-axis shows the negative logarithm of the adjusted p value. Point size represents the number of genes contributing to each term, and color indicates GO domain: Biological Process (pink), Cellular Component (green), or Molecular Function (blue). Selected GO terms associated with behavioral and neuronal processes are labeled for clarity.

QRP also had a significant, exposure-dependent effect on responsiveness to sucrose (94-96 individuals per group from each colony, 4 colonies; Figure 1B). Individuals in the LE group were significantly less responsive to sucrose than those in the other treatment groups (LE vs SE: z = -5.44, p < 0.0001; LE vs nE: z = -3.75, p = 0.00052). By contrast, individuals in the SE group did not differ significantly in sucrose responsiveness relative to the nE group (SE vs nE: z = 1.61, p = 0.241).

We repeated the same two experiments with individuals derived from queens that were naturally mated (NM); due to extreme polyandry, this results in greater genetic variation between individuals, and thus across treatment groups. Trends were similar to those for individuals derived from SDI queens (Figure S1A and B).

We next asked whether weakening social stimulation through stimulus cessation would also affect behavior over the same duration, leading to different outputs, as predicted by NTGT. We again exposed two groups of 3-day-old bees to QRP. In one group, QRP was removed after 10 min, whereas the other group remained exposed. After 1 hour, we then tested the two treated groups and the control group for retinue behavior, to measure their ability to respond to the QRP when QRP was present again (60 individuals per group from each colony, 6 colonies). After 1 hour of QRP treatment (SE), individuals showed greater interest in QRP than individuals in the control (nE) or 10 min groups (SE vs 10 min: z = 3.38, p = 0.002; SE vs nE: z = 4.57, p < 0.0001). The 10 min group showed a trend toward higher interest in QRP than the nE group, but this was not statistically significant (10 min vs nE: z = 2.21, p = 0.067) (Figure S1C). We also performed the same experiment with bees derived from NM queens. We again observed similar trends to those observed for individuals derived from SDI queens (Figure S1D).

These behavioral results are consistent with the predictions of NTGT and led us to test a key molecular prediction of the model.

### Experiment 2A: QRP stimulus strength differentially reshapes honey bee antennal lobe and mushroom body transcriptomes

The NTGT model predicts that gene expression is related to the strength of social stimuli; for QRP, this predicts that the weaker QRP stimulus induces smaller changes than a stronger stimulus. Bulk RNA sequencing (RNA-seq) was performed on two regions of the bee brain for individuals from the same three groups as in Experiment 1: SE (reflecting weaker social stimulus), LE (reflecting stronger social stimulus), and nE. The bees used for RNA-seq were from the same experimental cohort as those used in Experiment 1, collected before the behavioral assays were conducted.

The two brain regions we profiled with transcriptomics were the antennal lobes (AL) and the mushroom bodies (MB). The AL is the primary sensory neuropil for the perception and processing of olfactory information, including pheromones, and the MB is a major higher-order integration center involved with learning and memory and social responsiveness^22–24^.

Stronger social stimulus induced greater transcriptomic responses in both the AL and MB, whereas weaker stimulus induced a substantial response only in the AL. In the AL, LE individuals showed 2,593 DEGs and SE individuals showed 2,099 DEGs compared with the nE group, with 546 shared upregulated DEGs and 707 shared downregulated DEGs between the two treatment groups (Figure 1C, Table S1). In the MB, LE individuals showed 807 DEGs and SE individuals showed 79 DEGs compared with the nE group, with only 3 shared upregulated DEGs and 6 shared downregulated DEGs between the two treatment groups (Figure 1D, Table S1).

The genes ranked among the top DEGs differed with QRP stimulus strength in both brain regions. Genes differentially expressed in the SE group included immediate early genes, e.g., *hormone receptor 38* (*Hr38*), and other known fast response genes, e.g., *heat shock protein 83* (*Hsp83*). By contrast, genes differentially expressed in the LE group included downstream effectors of the juvenile hormone (JH) pathway, including *Krüppel homolog 1* (*Kr-h1*) (Table S1). QRP is known to affect long-term physiology and behavior in honey bees by modulating the JH signaling system^25,26^.

Gene Ontology (GO) analysis revealed distinct patterns of functional enrichment in the AL between the SE and LE groups. The neuronally related biological process terms uniquely enriched in the DEG list of the SE group were primarily associated with neurodevelopment, whereas those enriched in the DEG list of the LE group were related to neural circuit wiring (Figure 1E and F). The MB DEG lists showed substantially fewer significantly enriched GO terms than the AL DEG lists (Table S2).

### Experiment 2B: Individual variation in biological embedding reflects differences in cellular brain GRNs, as predicted by NTGT

In addition to the group trends reported above in the retinue assays of Experiment 1, we noted striking individual differences in behavioral responsiveness to QRP. This occurred despite the fact that all individuals had a high degree of genetic similarity and were reared in the same environment. Individual differences in social behavior are pervasive across animal and human societies and understanding their mechanistic basis is of great importance, from both biological and biomedical perspectives. In introducing NTGT, Traniello and Robinson (2021) suggested that individual differences in biological embedding can be explained by individual differences in the linkage between environmental responsiveness, neural activity, and brain gene expression^17^. Recent results for aggressive behavior indicate that these differences are related to differences in the topology of brain cell gene regulatory networks (GRNs), but the precise sensory inputs were not controlled in that study^27^. We performed single-nucleus brain GRN analysis to test this NTGT prediction as it relates to individual differences in QRP responsiveness.

We used single-nucleus RNA-sequencing (snRNAseq) to profile the transcriptomes of approximately 66,000 nuclei from the brains of HIGH-and LOW-QRP responding bees (N = 4 per group), identified in the retinue behavior assay from the bees exposed to shorter (SE) or longer (LE) periods of QRP stimulus (Figure 2A). We integrated all high-quality nuclei into a single dataset for direct comparison and projected the data into two dimensions using uniform manifold approximation and projection (UMAP) to visualize cellular heterogeneity. A total of 84 high-confidence cell clusters were identified (Figure 2B). Of these, 80 clusters were functionally annotated and categorized into 27 functional cell types using identified marker genes, together with reported markers from *Drosophila melanogaster* and published honey bee brain sc/snRNAseq atlases (Table S3).

**Figure 2.**
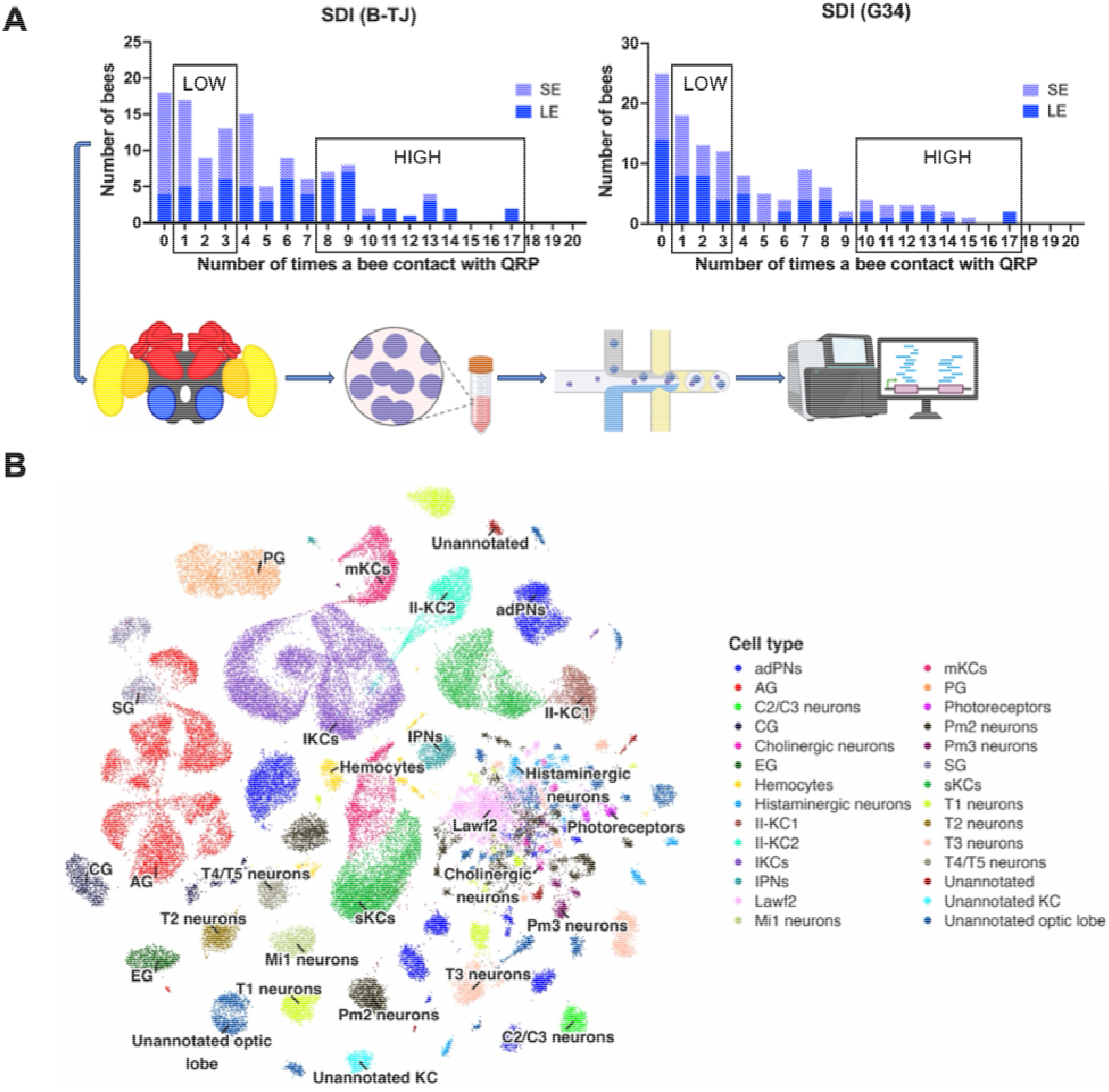
Individual honey bees showing differences in responsiveness to Queen Retinue Pheromone used to create brain single-nucleus transcriptomic profiles. **(A)** Selection of bees based on retinue response and experimental workflow for single-nucleus RNA sequencing (snRNAseq). The two colonies included in this experiment were also used in the earlier bulk RNA-seq analysis. During the 2024 retinue assays, the number of contacts with the QRP strip was recorded for each individual bee, and bees were ranked by contact frequency within each QRP exposure treatment. The 4 bees with the highest and the 4 with the lowest contact counts within each treatment were classified as high- and low-responsive (HIGH and LOW), respectively, and selected for snRNAseq. Brains were dissected from the selected bees, nuclei were isolated, and sn libraries were prepared for sequencing and downstream transcriptomic analysis. Histograms show the distributions of individual QRP contact frequencies for bees exposed to QRP for 1h (SE) or 3 days (LE), with the selected HIGH and LOW bees indicated. **(B)** Uniform manifold approximation and projection (UMAP) of the snRNAseq dataset showing transcriptionally defined nuclear clusters and their cell-type annotations. Each point represents an individual nucleus, and colors indicate the assigned cell type or neuronal subtype based on established marker-gene expression in *A. mellifera* and *D. melanogaster*. A total of 27 functional cell types were identified. Spatial proximity in the UMAP reflects similarity in transcriptional profiles and does not represent anatomical position within the brain.

We generated DEG lists for each cell type, comparing HIGH-and LOW-QRP responsive bees (responsiveness) in both LE and SE conditions (exposure). Across all exposure/responsiveness pairwise comparisons, cell clusters composed of Kenyon cells (KC) contained the largest number of DEGs (exposure: N = 1,313; responsiveness: N = 1,050), followed by optic lobe neuron clusters (OLN) (exposure: N = 1,092; responsiveness: N = 822), then glial clusters (exposure: N = 1,049; responsiveness: N = 725), and finally clusters from the olfactory projection neurons (OPN) (exposure: N = 377; responsiveness: N = 253) (Table S4).

Bulk RNA-seq of MB and AL (Experiment 2A) and snRNAseq revealed similar patterns of differential gene expression. In the MB, 286 DEGs were shared between the bulk and sn datasets (Fisher’s exact test: OR = 1.69, p = 1.54 × 10), with 75.5% showing the same direction of change (binomial test: 216/286, p < 2.2 × 10 ¹) and positively correlated effect sizes (Spearman’s: ρ = 0.44, p < 2.2 × 10 ¹). In the AL, 30 DEGs were shared (OR = 1.61, p = 0.016), of which 73.3% changed in the same direction (22/30; p = 0.016), with a stronger positive correlation in effect sizes (ρ = 0.68, p = 5.44 × 10) (Table S5).

To compare HIGH and LOW responders in both LE and SE conditions, we used the snRNAseq results to model cellular GRNs and Kolmogorov–Smirnov (KS) distance to compare regulatory dissimilarity^27,28^. This was done by integrating differences across cell-type-specific TF regulons, with each regulon comprising a transcription factor (TF) and its associated target genes (TGs) (Figure 3A). Overall, there was significantly greater GRN dissimilarity as a function of exposure compared to responsiveness (w = 940, p = 0.00026, Figure 3B). Examination of individual pairwise comparisons further showed that GRN dissimilarity between SE and LE individuals was more pronounced in the HIGH responders than the LOW responders (v = 1, p = 4.47 × 10, Figure 3C, left panel). In addition, GRN dissimilarity between LOW and HIGH bees was greater in response to long rather than short exposure (v = 0, p = 4.00 × 10, Figure 3C, right panel).

**Figure 3.**
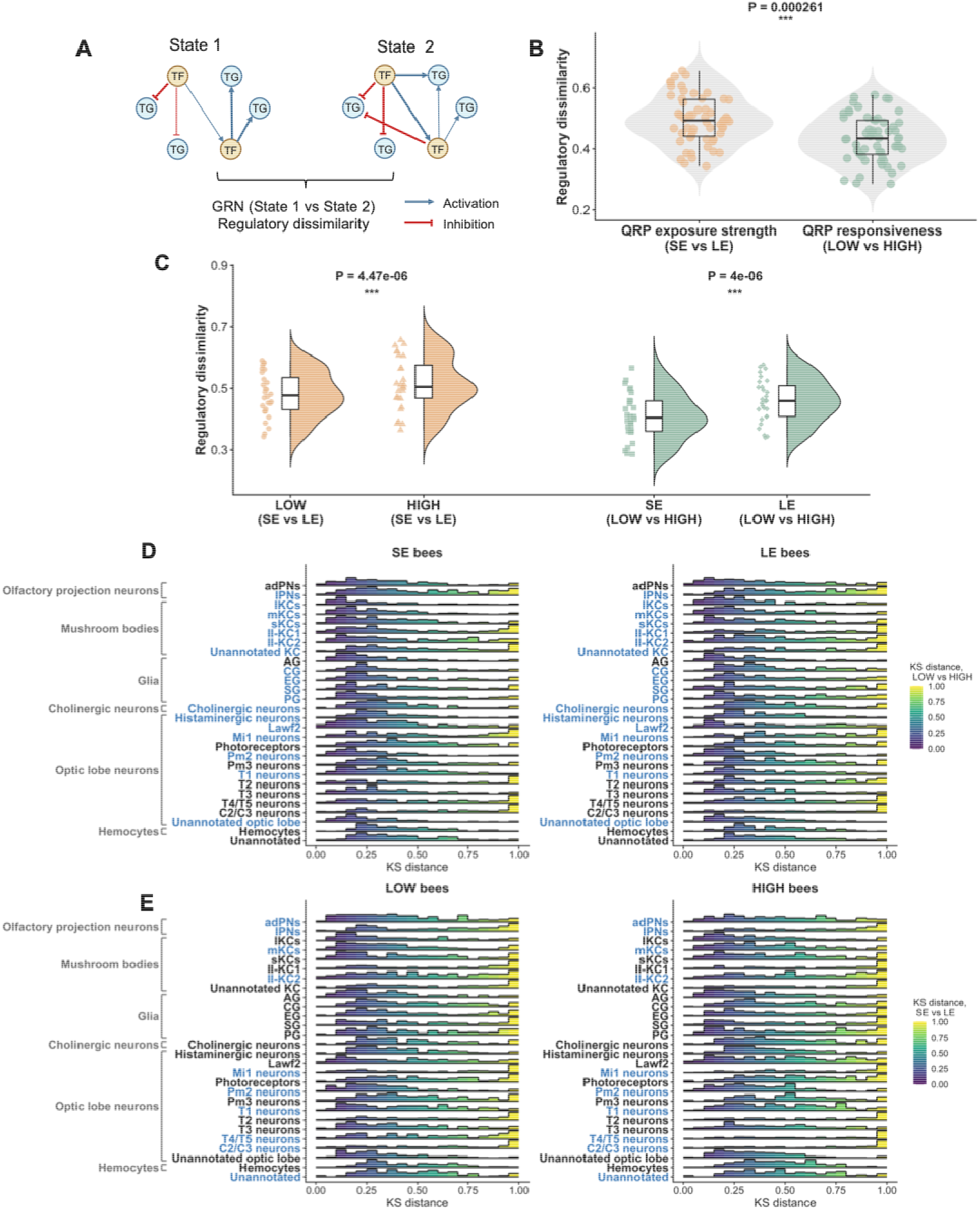
Individual differences in responsiveness to Queen Retinue Pheromone are associated with differences in the topology of brain cellular gene regulatory networks. **(A)** Schematic illustrating the comparison of gene regulatory network (GRN) states. Changes in predicted regulatory relationships between transcription factors (TFs) and target genes (TGs) were used to quantify regulatory dissimilarity between two states. **(B)** Overall regulatory dissimilarity associated with shorter or longer QRP stimulus exposure (SE and LE) and behavioral responsiveness to QRP (LOW and HIGH). Each point represents the Kolmogorov–Smirnov (KS) distance for one cell subcluster, with the distributions and boxplots shown for each comparison. Regulatory dissimilarity across cell subclusters was greater between SE and LE individuals than between LOW and HIGH individuals (***p = 0.000261). The KS distance was used to quantify GRN regulatory dissimilarity between two states, with larger values indicating greater divergence. QRP stimulus strength was varied as in Figure 1A. **(C)** Regulatory dissimilarity resolved according to behavioral response and QRP stimulus strength. Left, SE vs LE regulatory dissimilarity calculated separately for LOW and HIGH bees; the GRN dissimilarity between SE and LE was greater in HIGH-responsive bees compared with LOW-responsive bees (***p = 4.47 × 10 ^6^). Right, regulatory dissimilarity between LOW and HIGH individuals calculated separately within the SE and LE treatment groups; the GRN dissimilarity between LOW and HIGH bees was greater after the stronger QRP stimulus (3 days) than after the weaker QRP stimulus (1 h) (***p = 4.00 × 10 ^6^). Each point represents the KS distance for one cell subcluster, with density distributions and boxplots showing the corresponding distributions. **(D)** Cell-type-specific distributions of KS distances between LOW- and HIGH-responsive bees, shown separately for SE (left) and LE (right) bees. Blue-highlighted subcluster names indicate cell subclusters in which LOW vs HIGH regulatory dissimilarity differed significantly between SE and LE bees. An enlarged view with detailed annotations of selected cell types is provided in Figure S2A. **(E)** Cell-type-specific distributions of KS distances between SE and LE bees, shown separately for LOW-(left) and HIGH-responsive (right) bees. Blue-highlighted subcluster names indicate cell subclusters in which SE vs LE regulatory dissimilarity differed significantly between LOW- and HIGH-responsive bees. An enlarged view with detailed annotations of selected cell types is provided in Figure S2B. Ridgeline plots show the distributions of KS distances for each annotated cell subcluster. Each bar represents the proportion of TF regulons within a cell subcluster whose KS distances fall within the corresponding bin. Bar color indicates KS distance, with larger values (yellow) representing greater GRN regulatory dissimilarity. Cell subclusters are grouped by major brain cell classes, including olfactory projection neurons, mushroom body Kenyon cells, glia, cholinergic neurons, optic lobe neurons, and hemocytes. Statistical results are provided in Table S6.

Differences in GRN topological dissimilarity (LOW vs HIGH bees) between social stimulus strengths showed clear cell-type specificity and predicted functional specificity. These differences were strongest in glial cells and the KCs of the MB (Figure 3D, Figure S2A, Table S6). Overall, *lozenge* (*lz*) and *grain* (*grn*) were among the top-ranked TF regulons showing regulatory dissimilarity between LOW and HIGH bees in the KCs and glia, with the *lz* regulon showing greater dissimilarity in SE individuals and the *grn* regulon showing greater dissimilarity in LE individuals. Other TF regulons also showed significant cell-type-specific associations with GRN dissimilarity, including *mondo* and *ventral veins lacking* (*vvl*) in the MB, and *nuclear hormone receptor HR96* (*HR96*) and *nucleosome-destabilizing factor* (*Ndf*) in glia (Figure S3A and B).

Individual differences in GRN topological dissimilarity (SE vs LE bees) also were strong in sensory-processing cell types, including olfactory projection neurons (OPNs) and optic lobe neuron clusters (OLNs; Figure 3E, Figure S2B, Table S6). In the OPNs and OLNs, *clockwork orange* (*cwo*) and *traffic jam* (*tj*) were among the TF regulons showing the strongest differences associated with individual variation (Figure S3C and D).

These results further demonstrate that individual differences in behavior relate to individual differences in brain GRN topology, which provide the mechanisms to link cellular molecular states to neural states.

### Experiment 3: QRP treatment enhances antennal lobe neuronal activity

NTGT also predicts that exposure to strong stimuli that lead to large and sustained changes in brain gene expression causes changes in neuronal states and rewiring of neural circuits. We tested this prediction by examining whether longer exposure to QRP, which we have shown above to induce greater and more sustained changes in brain gene expression, also causes changes in electrophysiological activity. We used extracellular electrophysiological recordings to characterize AL neuronal responses following QRP exposure (Figure 4A). We chose the AL because it is the primary olfactory processing center where queen pheromones evoke neuronal responses^29^, whereas the MB exhibits sparse and more complex olfactory coding that would complicate the interpretation of extracellular recordings^30^. We recorded the extracellular voltage traces from LE and nE bees in response to QRP volatile exposure (Figure 4B).

**Figure 4.**
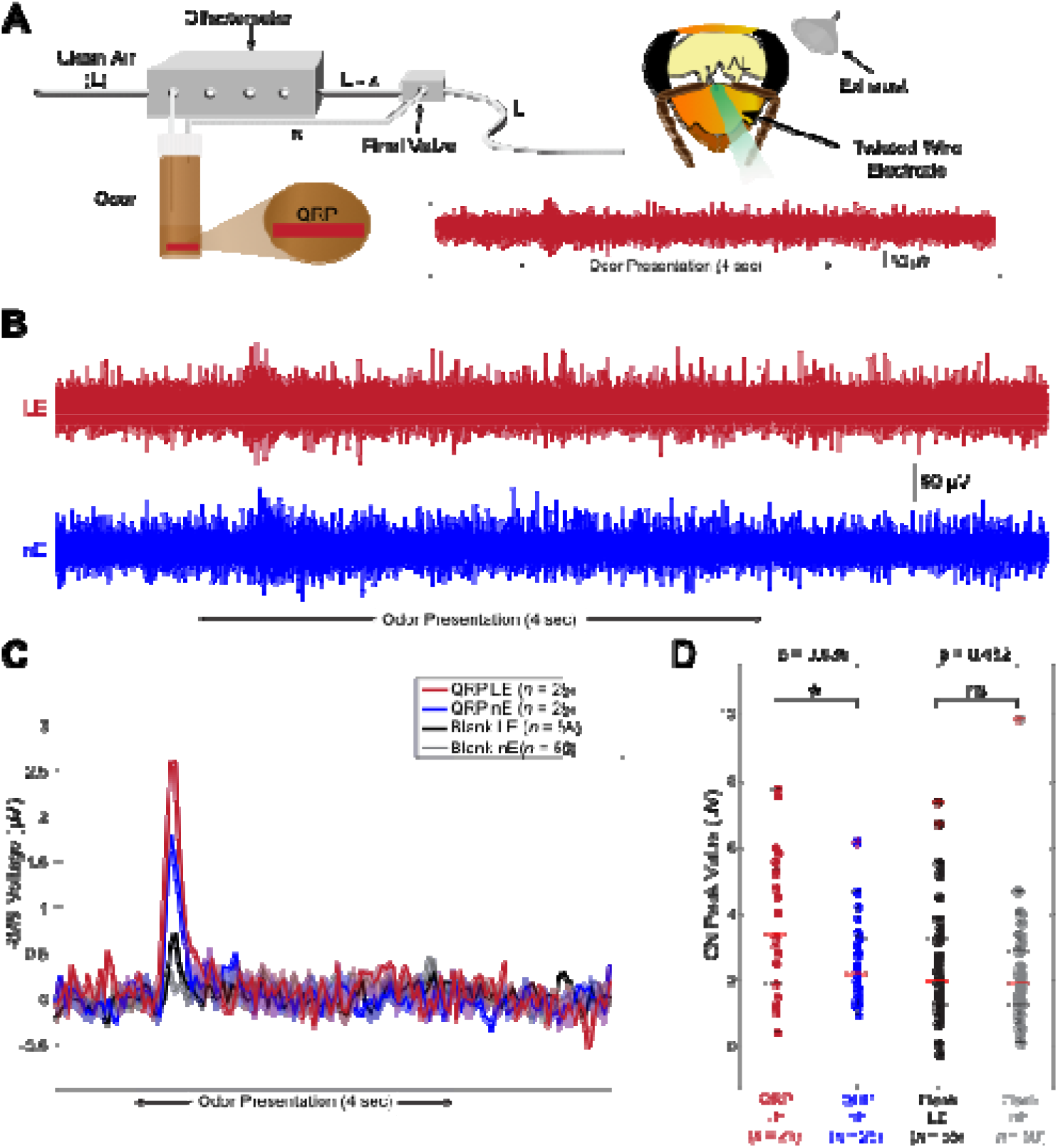
Exposure to Queen Retinue Pheromone caused changes in neuronal activity in the honey bee antennal lobe. **(A)** Schematic of the odor delivery and electrophysiological recording setup. A QRP strip was placed in an airtight vial, and a controlled airflow of clean air (L) was passed through the vial using an olfactometer to generate odor-laden air (L-x), which was delivered to the honey bee antenna, controlling odor volume and duration. Neural activity was recorded from the antennal lobe (AL) using a twisted-wire electrode, and residual odor was removed via an exhaust system. **(B)** Representative extracellular voltage traces recorded from QRP-exposed bees (3 days, LE) and control bees (nE) during QRP presentation. The shaded region indicates the 4 s odor presentation period. **(C)** Population root-mean-square (RMS) voltage responses to QRP and blank stimuli in LE and nE bees. For QRP stimulation, 25 trials were recorded from LE bees and 25 trials from nE bees. For blank stimulation, 55 trials were recorded from LE bees and 50 trials from nE bees. Lines show the population response over time, with shaded areas indicating the associated variation around the mean. The shaded background indicates the 4 s odor presentation period. **(D)** Comparison of ON peak response values across treatment and stimulus conditions. LE bees showed significantly greater responses to QRP stimulus than nE bees (*p = 0.036), whereas no significant difference was detected between LE and nE bees in response to the blank stimulus. Each point represents an individual odor trial. Boxes extend from the 25th percentile (Q1) to the 75th percentile (Q3), with the line within each box indicating the median. Whiskers extend to 1.5 times the interquartile range, and red crosses indicate outliers.

The population root-mean-square (RMS) voltage responses to QRP were larger in LE relative to nE individuals (Figure 4C). Consistent with this pattern, ON peak responses to QRP were significantly greater in LE relative to nE individuals (z = -2.10, p = 0.036). In contrast, ON peak responses to the blank stimulus (the clean odor) did not differ significantly between LE and nE individuals (z = -0.79, p = 0.43; Figure 4D). These results are consistent with the predictions of NTGT.

## Discussion

NTGT is a new conceptual framework for understanding the neural and molecular mechanisms underlying biological embedding to shape brain and behavior. Using honey bees and their highly stereotyped and time-dependent responses to QRP, we obtained behavioral, transcriptomic, and electrophysiological results consistent with NTGT. Our results also suggest that interactions between neural and molecular systems are shaped by individual differences in responsiveness to social stimuli, to enable individuals to uniquely anticipate future events. In this study we controlled one aspect of the environment, the strength of QRP stimulus (by varying duration of exposure); genetic and other environmental variables surely also influence differences in responsiveness.

QRP is a multi-component, multi-function pheromone central to social regulation in honey bee colonies. Our results are consistent with its known releaser and primer effects^20,21,31^; retinue behavior was affected by both weaker and stronger QRP stimuli, whereas sucrose responsiveness was affected only by the stronger QRP stimulus. Retinue behavior is a rapid releaser pheromone-type response mediated at the periphery and by AL neurons^32,33^. By contrast, sucrose response threshold is more of a primer pheromone-type response^21,34^. Our results indicate that releaser and primer effects are caused by differences in the intensity of the exposure to the pheromone itself and manifest differently in different regions of the brain; retinue behavior engaged genes in the AL while sucrose response threshold engaged genes in both AL and MB. These graded responses are reminiscent of observations of the responses to stimulus intensity in human pain perception^35^. Our results are consistent with the NTGT prediction that stronger social stimuli are more likely to shape future behavior.

As noted above, QRP stimulation induced spatiotemporally regulated transcriptional changes in the honey bee brain, with gene expression patterns differing between brain regions and across stimulus strengths. Stronger exposure to QRP induced broad transcriptional changes across the sucrose responsiveness pathway, affecting genes that are predicted to reshape reward-related circuits, whereas weaker exposure to QRP produced little pathway-level remodeling, leaving sucrose responsiveness unchanged. In the ALs, by contrast, weaker (1-hour exposure) and stronger QRP stimuli (3-day exposure) elicited highly similar expression changes, which may explain why retinue behavior was altered after both stimulus strengths. AL networks may therefore remodel more rapidly than networks in higher-order brain regions, allowing behaviors that depend on peripheral sensory processing to shift faster, as in honey bee aggression, which can change within minutes of encountering an intruder^36^. This is consistent with evidence that some behaviors are adjusted locally within sensorimotor circuits^37^.

We observed strong variation in QRP responsiveness among closely related individuals, consistent with widespread observations of individual differences in responsiveness to environmental stimuli in vertebrates, including humans^3,38,39^. Pre-existing differences in neuronal network states can predispose individuals to respond differently to the same challenge^40^. Perhaps this explains why we observed that QRP exposure strength had a stronger effect on the topology of brain GRNs in cells of HIGH-responsive bees, with GRN dissimilarity between LOW- and HIGH-responsive bees being greater after the stronger QRP stimulus than after the weaker QRP stimulus. The stronger QRP stimulus may recruit additional transcriptional programs beyond those induced by a brief stimulus^41^, with the resulting changes in GRN activity acting on neuronal networks to influence subsequent responsiveness^42^.

QRP-induced GRN changes showed strong cell-type specificity, with different bioinformatically defined clusters associated with differences in QRP stimulus strength and responsiveness. A stronger QRP stimulus affected more cell clusters than did a weaker QRP stimulus. Moreover, glial cells showed pronounced, strength-dependent GRN dissimilarity between HIGH- and LOW-responsive bees, suggesting that glial cells may act as a signaling relay converting short-term exposure into long-term effects. A comparable role has been described in mice, where astrocytes serve as a critical relay in a stress-induced positive feedback loop in the lateral habenula–locus coeruleus axis^43^. In glia, the *lz* regulon showed the greatest dissimilarity between HIGH- and LOW-responsive bees after one hour of exposure and the *grn* regulon after three days. Given the roles of *lz* in upstream cell-fate regulation^44^ and *grn* in motor axon guidance^45^, weaker and stronger QRP stimuli may affect different stages of neural regulatory processes.

At the cellular level, brain GRN dissimilarity for bees experiencing either a weaker or stronger QRP stimulus showed spatial differences in HIGH- and LOW-responsive bees. It was concentrated in sensory-processing cell types such as AL projection neurons, which process queen mandibular pheromone in honey bees^29^, possibly reflecting a role of the AL in modulating responses to social cues^46^. Previous GRN analyses have linked the TF *cwo* to individual variation in egg-laying and foraging behavior, consistent with the pattern reported here^47^. Our findings show how NTGT can provide a heuristic framework for understanding individual differences in behavior.

Our electrophysiological data support the prediction that neural circuit rewiring occurs after changes in brain gene expression to shape future behavior. AL electrophysiological responses to QRP were stronger in bees previously exposed to a stronger QRP stimulus, suggesting that prior experience had modified circuit function rather than merely engaging the circuit at the time of recording. Experience-dependent enhancement of neural responsiveness may be a conserved feature of sensory systems. In the noctuid moth *Spodoptera littoralis*, for example, brief pre-exposure to sex pheromone potentiates both behavioral responses and AL neuron activity to the same pheromone^48^. A parallel phenomenon has been described in mammals, where olfactory fear conditioning increases the number of odor-specific sensory neurons and enlarges the corresponding olfactory glomeruli, changes that can be reversed by extinction training^49^.

Our study touches on the relationship between instinct and learning. Learning is defined as a change in behavior due to experience^50^. Biological embedding also involves experience-dependent changes in behavior, for both instinctive and learned behaviors. Instincts and learned behaviors are hypothesized to share overlapping molecular substrates^19,51–53^, with neural plasticity unfolding across multiple timescales^54,55^. These findings suggest that the NTGT framework can be applied to both instinctive and learned behaviors, with relevance to the evolution of emotions in humans. Recent research on the mechanisms underlying models of emotion has highlighted specific patterns of neural activity in brain circuits^56^; it is not difficult to envision how NTGT could be used in the future to study neural and molecular bases of emotion.

## Materials and Methods

### Experimental subjects

We used two types of honey bee colonies as sources of bees for our experiments: colonies derived from queens each instrumentally inseminated with a single drone (SDI colonies), and colonies derived from queens that were naturally mated (NM colonies). Queens naturally mate with 15-20 males, thus individuals from an SDI colony are highly related to each other (average relatedness = 0.75 due to haplodiploidy) compared with individuals from an NM colony. In 2024, we used 4 SDI and 3 NM colonies for the sucrose responsiveness and retinue assays, and 2 SDI colonies for the bulk RNA-sequencing (RNA-seq) and single-nucleus RNA-seq (snRNAseq) experiments. In 2025, we used 6 SDI and 6 NM colonies for the retinue behavior assays. All colonies were maintained according to standard beekeeping practices at the University of Illinois Bee Research Facility, Urbana, Illinois, USA. Colonies in this area are a mixture of European subspecies of *Apis mellifera*, primarily *A. mellifera ligustica*.

### Pheromone treatment

Two days before adult eclosion, we removed honeycomb frames containing pupae from SDI and NM colonies and maintained them in a dark incubator at 34 °C and 50% relative humidity. We collected one-day-old newly emerged adult individuals for pheromone treatment. They were exposed to the nine-component synthetic queen retinue pheromone (QRP), embedded in a plastic strip (Intko Supply Ltd., Chilliwack, BC, Canada) for different durations. QRP strips were cut into 1 cm lengths, each containing approximately two queen equivalents^32^. An untreated strip of the same color and dimensions was used as a control.

Bees were collected from each colony, individually marked with colored numbered tags, and placed in Plexiglas cages (10 × 10 × 10 cm) at 50 bees per cage (bees in each cage were from the same colony). 6 cages were prepared and randomly assigned to 3 treatment groups, with 2 cages per group.

In the 2024 section of Experiment 1, in one treatment group, bees were continuously exposed to QRP strips for 3 days (a longer exposure, LE), representing a stronger social stimulus. In another treatment group, bees were kept without QRP for 3 days and then exposed to QRP for 1 hour (a shorter exposure, SE), representing a weaker social stimulus. Control bees were never exposed to QRP during the 3-day period (Untreated, nE). All boxes were supplied with pollen paste, honey, and drinking water that were available at all times, and were kept in a dark incubator at 34 °C and 50% relative humidity. After 3 days of treatment, bees from each cage were used either for behavior or frozen for RNA-seq analysis. One subset was flash-frozen in liquid nitrogen immediately before the behavioral assays and stored at -80 °C while the second subset was used for sucrose response threshold or retinue analysis and was flash-frozen immediately after testing. A total of 4 SDI colonies and 3 NM colonies were used, for a total of 3,548 individuals. Bees were sampled and tested in the behavioral assays at the same time of day.

Unlike in 2024, bees were not exposed to QRP in the 3 days prior to the experimental treatments. Rather, in the 2025 section of Experiment 1, some individuals were exposed to QRP strips for 1 hour (SE), while others were exposed to QRP strips for 10 min, after which the QRP strips were removed and replaced with control strips for the remaining 50 min (10 min). This design allowed us to test whether weakening social stimulation through stimulus cessation, within the same 1-hour period, would lead to different behavioral outcomes, as predicted by NTGT. Control bees were never exposed to QRP during the 1-hour period (nE). After treatment, they were used in retinue assays. Bees were caged and maintained as described above. 6 cages (50 bees each) were randomly assigned to the 3 treatment groups, with 2 cages per group. A total of 6 SDI colonies and 6 NM colonies were used, for a total of 2,340 individuals.

### Behavior assays

#### Sucrose response threshold analysis

We used the standard proboscis extension reflex (PER) testing method with slight modifications^21,31,57^. Simple restraining harnesses for individual honey bees were made from plastic soda straws. Test bees were chilled on ice until they stopped moving. Once an individual was immobile, it was removed from the ice to avoid overexposure to cold and placed in a restraining harness. It was positioned in the harness with the dorsal thorax facing the cut-away portion of the straw, the head just above the top of the tubing, and the mouthparts extending beyond the edge of the straw, with the forelegs free. For each treatment, approx. 48 bees were fixed individually in plastic straws, leaving their antennae, proboscis, and forelegs free. Bees were then acclimated to ambient conditions (approximately 28 °C) for about 1 hour. After acclimatization, each bee’s antennae were touched with water. If the bee showed proboscis extension, it was allowed to drink water to satiation.

Sucrose solutions at concentrations of 0%, 10%, 20%, 30%, 40%, and 50% (w/v) were prepared using molecular biology grade water (Thermo Fisher, USA). A small droplet of each sucrose solution was delivered with a 10 µL pipette tip and gently touched to the antenna of each bee for approximately 5 seconds. Solutions were presented sequentially in ascending order of concentration. Proboscis extension responses (scored as 1 for extension and 0 for no extension) were recorded for each concentration. An inter-trial interval of 5 min was used between tests of different sucrose concentrations to reduce potential sensitization effects.

#### Retinue behavior

We used a previously described retinue assay to test the response of QRP-treated bees^20,32^. 15 individual bees were gently transferred from the treatment cages into 150 × 25 mm Petri dishes under red light to avoid inducing the release of alarm pheromones. After a 1 cm QRP strip was placed in the middle of dish, bees contacting the strip were recorded 30 s later and every 30 s thereafter for 10 min. Only contacts of the strip with mouthparts or antennae were recorded. A total of 20 counts were recorded. Strips were used only once, and groups of bees were immediately preserved in liquid nitrogen after testing. Each treatment included 4 replicate dishes.

For snRNAseq analyses of individual differences in responsiveness to QRP, we recorded the number of times each individual contacted the QRP strip during the retinue assays in 2024 and ranked bees by contact frequency. Within each treatment group, the 4 bees with the highest and the 4 with the lowest contact counts were selected for sequencing.

### Bulk RNA sequencing of antennal lobes and mushroom bodies

For Experiment 2A, performed in 2024, we used QRP-treated bees that were collected before behavioral assays were performed. The heads of individual bees were removed, and the cuticle around the brain, except for the eyes, was carefully opened in a dry ice and ethanol bath. The prepared heads were then transferred to prechilled RNA*later*-ICE solution (Thermo Fisher, USA) and stored at -20 °C for 16 h. After incubation in RNA*later*-ICE, brains were fully dissected on wet ice, and the upper part of the midbrain containing mostly the mushroom bodies (MBs) and antennal lobes (ALs), was carefully removed and stored at -80 °C until RNA extraction, as previously described^36,58^. Total RNA was isolated from MB and AL using the RNeasy Mini Kit (QIAGEN, Germany) with DNase treatment (Invitrogen, Carlsbad, CA, USA) and finally resuspended in 30 μL of RNase-free water. RNA concentration was determined using a NanoDrop 1000 spectrophotometer (Thermo Fisher Scientific, USA) and a Qubit 2.0 fluorometer (Thermo Fisher Scientific, USA), and RNA quality was assessed using a Bioanalyzer (Agilent Technologies, USA). RNA sequencing libraries were prepared with sequencing kits V1.0 (Illumina, USA); libraries were pooled, quantified via qPCR, sequenced on a single 25B lane of a NovaSeq X Plus instrument (Illumina, USA), generating 151 cycles from both ends of the fragment. Fastq files were produced and demultiplexed with the bcl-convert conversion software (version 4.1.7, Illumina, USA), and the resulting reads were 150 nucleotides in length. We sampled 2 SDI colonies across 3 treatment groups. 10 bees per colony per treatment were collected, and 2 tissues per bee were sequenced, yielding 120 sequencing libraries in total (2 × 3 × 10 × 2).

### Single-nucleus RNA sequencing (snRNAseq)

Before brain dissection, we screened the bees for viruses to ensure that the individuals used for snRNAseq were virus-negative. Brains were then dissected in a bath of dry ice and ethanol, and glands, tracheae and any surrounding non-brain tissues were removed. For each sample, we pooled 4 frozen brains from bees with the same QRP contact frequency category (HIGH-responsiveness or LOW-responsiveness) into a single tube. For each QRP treatment (LE and SE), we collected both HIGH and LOW bees from each colony. In total, this generated 8 pooled brain samples from 2 SDI colonies; the same colonies were also used for bulk RNA-seq.

Samples were processed following the nucleus isolation protocol using the Chromium Nuclei Isolation Kit (10x Genomics, USA). Briefly, 200 µL of 0.25× lysis buffer was added to the tube, and the brain was dissociated with a pestle on ice. Once no visible unhomogenized brain tissue remained, an additional 300 µL of 0.25× lysis buffer was added. The sample was mixed by pipetting 10 times and incubated on ice for 10 min. The lysate was then filtered through a pre-chilled nuclei isolation column into a 2 mL collection tube by centrifugation at 16,000 × g for 20 s at 4 °C to collect the nuclei. The nuclei were resuspended by vortexing for 10 s at maximum speed, followed by centrifugation for 6 min at 500 × g at 4 °C to pellet the nuclei. The supernatant was discarded, and the nuclei pellet was resuspended in 500 µL debris removal buffer and gently mixed by pipetting until no pellet was visible. The suspension was centrifuged at 700 × g for 10 min at 4 °C to pellet the nuclei. The supernatant was discarded, and the nuclei pellet was resuspended in 1000 µL wash and resuspension buffer containing RNase inhibitor, followed by centrifugation at 300 × g for 10 min at 4 °C to pellet the nuclei. Finally, the supernatant was discarded, and 200 µL wash and resuspension buffer was added to resuspend the nuclei for quality control and library preparation.

Nucleus suspensions were kept on ice and transported immediately to the Roy J. Carver Biotechnology Center at UIUC. Nucleus concentration was quantified using an AO/PI staining based total count assay on a Nexcelom Cellometer Spectrum image cytometry system (Nexcelom Biosciences, USA). Nucleus integrity was then assessed by microscopic inspection to confirm that the majority of nuclei within the field of view were intact and displayed normal morphology. Libraries were constructed using the chromium GEM-X single cell 3’ reagent kits v4 (10x Genomics, USA). The pooled libraries were quantified by qPCR and sequenced on two NovaSeq X Plus 25B lanes using the sequencing kit v1.0 (Illumina, USA), with 28 bp for read 1 and 150 bp for read 2. FASTQ files were generated and demultiplexed.

### Bioinformatics

#### Bulk RNA-seq

Approximately 30 million cleaned paired-end reads were obtained per sample. Read quality was assessed using FastQC v0.11.9^59^. Reads were aligned with STAR v2.7.10a^60^ using default parameters to the honey bee reference genome assembly HAv3.1^61^ and to reference genomes of the following honey bee viruses: Deformed wing virus (DWV, GCF_000852585.1), Varroa destructor virus 1 (VDV, GCF_000856945.1), Acute bee paralysis virus (ABPV, GCF_000856345.1), Israeli acute paralysis virus (IAPV, GCF_000870485.1), Sacbrood virus (SBV, GCF_000847625.1), and Chronic bee paralysis virus (CBPV, GCF_000875145.1). Reads were first aligned to the viral reference genomes, and unmapped reads were subsequently aligned to the honey bee reference genome. Read counts were generated using the featureCounts function in the Subread package v2.0.4 with default parameters^62^.

Prior to differential gene expression analysis, lowly expressed genes were removed; genes were retained only if they had at least 1 count per million (CPM) in at least as many samples as the smallest group size. Subsequently, we determined differential gene expression between the 3 treatments (LE, SE, and nE) by 3 pairwise comparisons (LE vs nE, SE vs nE, and LE vs SE) for 2 tissues (AL and MB) using the limma-trend v3.62.2 accordingly^63^.

In both MB and AL, a total of 13 samples showed varying levels of reads mapping to a common honey bee virus. To control for variation in viral load across samples, viral load was included as a covariate in the linear model. In addition, because samples originated from different colonies, colony was treated as a blocking factor.

We then determined the number of differentially expressed genes (DEGs) for each comparison using the adjusted p value (FDR) < 0.05. For Gene Ontology (GO) enrichment analysis, we first converted *A. mellifera* genes to *Drosophila melanogaster* (GCF_000001215.4_Release_6_plus_ISO1_MT) orthologs using three methods: RBH-BLAST^27^, BLASTp^64^, and gProfiler2^65^. We combined the results from all three methods to generate one-to-one ortholog pairs, and used this final ortholog list to convert the DEGs to *D. melanogaster* genes. GO over-representation analysis was performed with the enrichGO function in the R package clusterProfiler v4.14.6 using org.Dm.eg.db, with Benjamini-Hochberg correction and significance thresholds of adjusted p < 0.05^66^.

#### Single-nucleus RNA sequencing (snRNAseq)

snRNAseq reads were processed using Cell Ranger v9.0.1 for alignment and quantification. Alignment was performed against an integrated reference containing the honey bee genome and the common honey bee virus genomes described above. Quality control and data processing were performed in Seurat v5^67^. First, nuclei were retained only if they had at least 1,180 detected genes (nFeature_RNA). This cutoff was defined using the highest-quality sample, where the minimum observed value was 1,180, and was then applied to all samples. In addition, nuclei with a high proportion of mitochondrial transcripts (defined as values more than three median absolute deviations above the median) were removed. After filtering, the maximum mitochondrial transcript proportion per nucleus was 0.42%, and no reads mapping to the common honey bee viruses were detected in any sample. Following quality control filtering, snRNAseq profiles for approximately 6,500 to 9,000 high-quality nuclei per sample were retained for downstream analyses.

Data were normalized using SCTransform (method = “glmGamPoi”, vst.flavor = “v2”), retaining 2,000 variable features ranked by residual variance for downstream analyses.

We integrated all nuclei according to sample ID by using Harmony v0.1.0^68^. To determine an appropriate clustering resolution for the 66,466 nuclei, graph-based unsupervised clustering was performed using FindNeighbors and FindClusters across a range of resolution values (0.1-1.0, in 0.1 increments). The relationships among clustering solutions at different resolutions were visualized using clustering trees generated with clustree v0.5.1^69^. Based on the clustree output, a resolution of 0.7 was selected for downstream analyses. We extracted the Harmony-corrected low-dimensional embeddings from the Seurat object and selected dimensions 1-40, which together explained 90% of the total variance. Uniform manifold approximation and projection (UMAP) visualizations were generated using these 40 harmony dimensions, with min.dist set to 0.75 to more evenly distribute nuclei in the embedding. Parameter choices were guided by previous honey bee sc/snRNAseq studies^27,70–72^ and were further adjusted to reflect the large number of nuclei recovered in our experiment and our experimental design.

Using these settings, we identified a total of 84 nucleus clusters. We identified conserved marker genes for each cluster using the ‘FindConservedMarkers’ function of Seurat. Significance was defined based on the Wilcoxon rank-sum test (combined p value < 0.05, log_2_FC > 0.5). Only genes detected in at least 10% of the nuclei within the cluster were considered. We then annotated nucleus types by matching cluster marker genes to published markers from *D. melanogaster*^73,74^ and *A.* mellifera^27,70,71,75^.

Within each cluster, differential expression was assessed using MAST with default settings^76^, with DEGs defined as genes with an adjusted p value < 0.05, for the contrasts LE vs SE and HIGH vs LOW. In addition, we compared DEGs from the snRNAseq analysis with the DEG sets identified in the bulk RNA-seq analysis for LE vs SE bees. Comparisons were restricted to genes tested in both datasets. Enrichment of shared DEGs was assessed using a one-sided Fisher’s exact test based on the hypergeometric distribution, testing whether the observed overlap exceeded that expected under independence. For genes identified as DEGs in both datasets, directional concordance was assessed by comparing the signs of the differential-expression effect estimates from the two datasets. The proportion of genes showing concordant directions of change was tested against a null expectation of 0.5 using a binomial test. Concordance in effect size was additionally assessed using Spearman’s rank correlation.

#### Gene regulatory network (GRN) analysis

We used SimiC for our single-nucleus gene regulatory network (GRN) analysis. SimiC infers phenotype-specific GRNs while imposing a similarity constraint across conditions, which allows direct comparison of regulatory architecture between experimental groups^28,77^. SimiC infers transcription factor (TF)-target gene (TG) relationships using imputed scRNAseq expression profiles to estimate regulatory interactions between each TF and its predicted target genes, with the set of genes predicted to be regulated by each TF constituting its inferred regulon. This approach has been successfully applied to honey bee brain scRNAseq data to compare differences in brain GRNs across other behavioral phenotypes^27^.

We downloaded the D. melanogaster TF list from TFLink78 and retained only TFs with biochemical and/or genetic validation. These TFs were then mapped to *A. mellifera* using a robust *A. mellifera*-*D. melanogaster* ortholog list, yielding 321 honey bee TFs as input for SimiC (Table S7). We used a curated list of honey bee TFs to model regulatory relationships with 2,000 TGs. Separate TF-by-TG incidence matrices were reconstructed for the four experimental phenotypes: LE_HIGH, LE_LOW, SE_HIGH, and SE_LOW. The joint inference was performed using regularization parameters set to lambda_1 = 0.1 and lambda_2 = 0.1, ensuring robust optimization across states while capturing phenotype-specific regulatory activity. The Regulon Activity Score, defined as the collective influence of a TF and its predicted TGs, was quantified using a weighted area under the curve scoring pipeline. For each individual cell, the regulon activity score measured the relative activity of a given regulon weighted by the expression levels of its associated TGs. Consequently, elevated regulon activity score values indicated a high regulatory influence of the TF on a cell’s transcriptional profile.

We calculated regulon activity score values independently for every cell across all identified clusters. To quantify the regulatory dissimilarity between phenotypes, we applied the Kolmogorov-Smirnov (KS) distance to the regulon activity score distributions within each cell type. This metric allowed for the identification of TFs exhibiting significant shifts in regulatory activity across behavioral states. We used a Wilcoxon rank-sum test to assess differences in GRN dissimilarity within each functional cell type.

### Electrophysiological recording in the honey bee antennal lobe

#### Surgery

Honey bees were collected individually into plastic vials and briefly anesthetized on ice until immobilized. Each bee was then secured in a custom-made 3D printed plastic restraining harness using dental wax (Surgident Periphery Utility Wax), applied to the posterior of the head to restrict head movement while allowing free movement of the antennae and mouthparts. Following harnessing, bees were left undisturbed at room temperature for 30-60 min, then fed 50% sucrose solution to satiation. Subsequently, QRP-treated bees were housed with a QRP strip and remained exposed to QRP during the overnight harnessing period, so that total exposure was 3 days, exactly as in the LE group. The control bees were housed in a separate cabinet to prevent contamination of QRP from the strip within the treatment group’s box. The following morning, feeding motivation was assessed as an indicator of physiological viability for electrophysiological recordings. Bees were presented with 50% sucrose solution to the antennae, and individuals that exhibited a proboscis extension response were used in electrophysiological experiments. To prepare for recording, antennae were immobilized using eicosane wax (Sigma-Aldrich) applied to the base of the scapes and pedicels and positioned with both flagella oriented anteriorly. The head capsule was then carefully shaved using a microscalpel, and bees were allowed to rest undisturbed for 10-20 min. A small window was subsequently cut in the head capsule between the antennae and below the ocelli. Glands and tracheae were removed to expose the brain, which was continuously perfused with a Ringer’s solution (in mM: 37 NaCl, 2.7 KCl, 8 Na2HPO4, 1.4 KH2PO4, pH 7.2; chemicals from Sigma-Aldrich) as described by previous literature^79,80^. The AL was desheathed immediately prior to electrode placement.

#### Odor vial preparation and presentation

QRP was placed into individual vials, and an empty vial was used as a blank control. Odor stimuli were delivered using a previously established protocol^80–84^. A commercial olfactometer (Aurora Scientific, 220A) was used to ensure precise and reproducible odor presentation. At the start of each trial set, 200 standard cubic centimeters per min (sccm) of zero-contaminant air was directed through the stimulus flow line (1/16 in. PTFE tubing) toward the honey bee antenna, while an additional 200 sccm was directed through a parallel dilution flow line to the exhaust. The outlet of the stimulus line was positioned approximately 2-3 mm from the distal antennal segment. Five seconds prior to stimulus onset, 40% of the dilution flow (80 sccm) was redirected through the odor line upstream of the odorant source (odorant vials). The odorized air then recombined downstream with the remaining 120 sccm of clean dilution air, ensuring complete mixing before reaching the final valve. This mixture was routed to the exhaust to allow the system to equilibrate and prime the line with odorized air. At stimulus onset, the final valve redirected the odorized air mixture through the stimulus line to the antenna, while the clean air stream was diverted to the exhaust. Odor delivery was maintained for 4 seconds at a constant total flow rate. Following stimulus offset, the final valve returned clean air to the stimulus line and redirected the odorized flow back to the exhaust. One second later, the 80 sccm odor flow was returned to the dilution line to prevent depletion of odorant headspace. This design maintained a constant total flow rate throughout the experiment, minimizing mechanosensory artifacts associated with fluctuations in air pressure. To facilitate rapid odor clearance, a 6-inch diameter funnel connected to a low vacuum source was positioned directly behind the bee during odor presentation.

#### Electrophysiology

*In vivo* extracellular recordings were obtained from honey bees exposed to QRP and blank controls. Two groups were tested: LE and nE bees. For odor-evoked responses to the QRP-containing vial, data were collected from 5 LE bees (25 trials total) and 5 nE bees (25 trials total). When recording odor-evoked responses from the blank vial, neural data were collected from 11 LE bees (55 trials total) and 10 nE bees (50 trials total). Each recording trial lasted 20 seconds, consisting of a 10-second baseline period, 4 seconds of odor stimulus, and a 6-second post-stimulus period. The trials were repeated 5 times with a 60-second inter-stimulus interval, yielding 5 trials of neural response data per odor for each individual.

Following surgical preparation, a silver chloride ground wire was placed in the Ringer’s solution within the head capsule. Neural activity from antennal lobe neurons was recorded using a custom-built 4-channel twisted-wire electrode (impedance: 350-450 kΩ) inserted superficially into the center of the antennal lobe. Voltage signals were sampled at 20 kHz and digitized using an Intan RHD 16-channel head stage (C3334). Digitized signals were transmitted to the Intan RHD interface board (C3100) and recorded using the Intan GUI in conjunction with LabVIEW for data acquisition and storage.

#### RMS transformation of neural voltage response

Voltage recordings were imported into MATLAB and bandpass filtered at 300 Hz before further analysis. For RMS processing, large-amplitude, broad voltage deflections associated with palp movements were identified as signals exceeding 11 standard deviations from the mean voltage amplitude. These movement artifacts were removed by replacing 80-time bins (4 ms) centered on each peak with the mean voltage value. For the QRP, a total of 100 voltage traces were analyzed (5 tetrodes × 2 odors × 5 trials per group, across treated and control groups). Only 1 trace from the treated group contained movement artifacts, while no artifacts were observed in control recordings. For the blank odor, 105 traces were analyzed (treated: 11 tetrodes × 1 odor × 5 trials = 55; control: 10 tetrodes × 1 odor × 5 trials = 50), of which 3 treated and 1 control traces contained movement artifacts. Filtered data were then trimmed to the time window of interest and processed using an RMS approach as described previously^80,81,83,84^. Briefly, a continuously moving 500-point RMS filter followed by a 500-point moving average filter was applied to the voltage traces. Baseline activity, defined as the mean signal during the 2 seconds preceding stimulus onset, was subtracted to obtain ΔRMS values. These values were subsequently binned into non-overlapping 50 ms intervals and the mean of each bin was calculated.

### Statistical analyses

Each individual was tested repeatedly in behavioral assays, so observations within individuals were not independent. In the PER assay, responses across successive solutions (water and increasing sucrose concentrations) are serially correlated within the same bee. Similarly, in the retinue response assay, each bee was observed repeatedly across 20 consecutive counts for contacts with the QRP strip. Therefore, we used generalized estimating equations (GEE) implemented in R (geepack::geeglm) to analyze main effects and interactions while accounting for within-individual correlation.

For PER, we fitted a GEE model with a binomial distribution and a logit link to relate the probability of a positive response to sucrose concentration and treatment. We specified a first-order autoregressive working correlation structure to account for serial correlation across repeated measurements within individuals.

For the retinue response assay, the outcome was a count (number of contacts). We fitted a GEE model with a Poisson distribution and a log link, using the form Number ∼ Time + I(Time^2) * treatment. Because the effect of Time was nonlinear and differed by treatment, we included the interaction between treatment and the quadratic term I(Time^2). The working correlation structure was again specified as the first-order autoregressive correlation structure.

For the AL electrophysiological data, each neuron had an RMS odor-evoked response to each odor for 5 different trials. In the statistical analyses, the trials were kept separate instead of being averaged together to avoid losing any important information or variability. The peak value was found for every trial and sorted into specific odors. An unpaired Wilcoxon rank-sum (Mann–Whitney U) test was performed comparing the peak response values of all neurons and trials of LE vs nE bees for the same odors (LE response to blank compared to nE response to blank, LE response to QRP compared to nE response to QRP).

---

**Figure S1.**
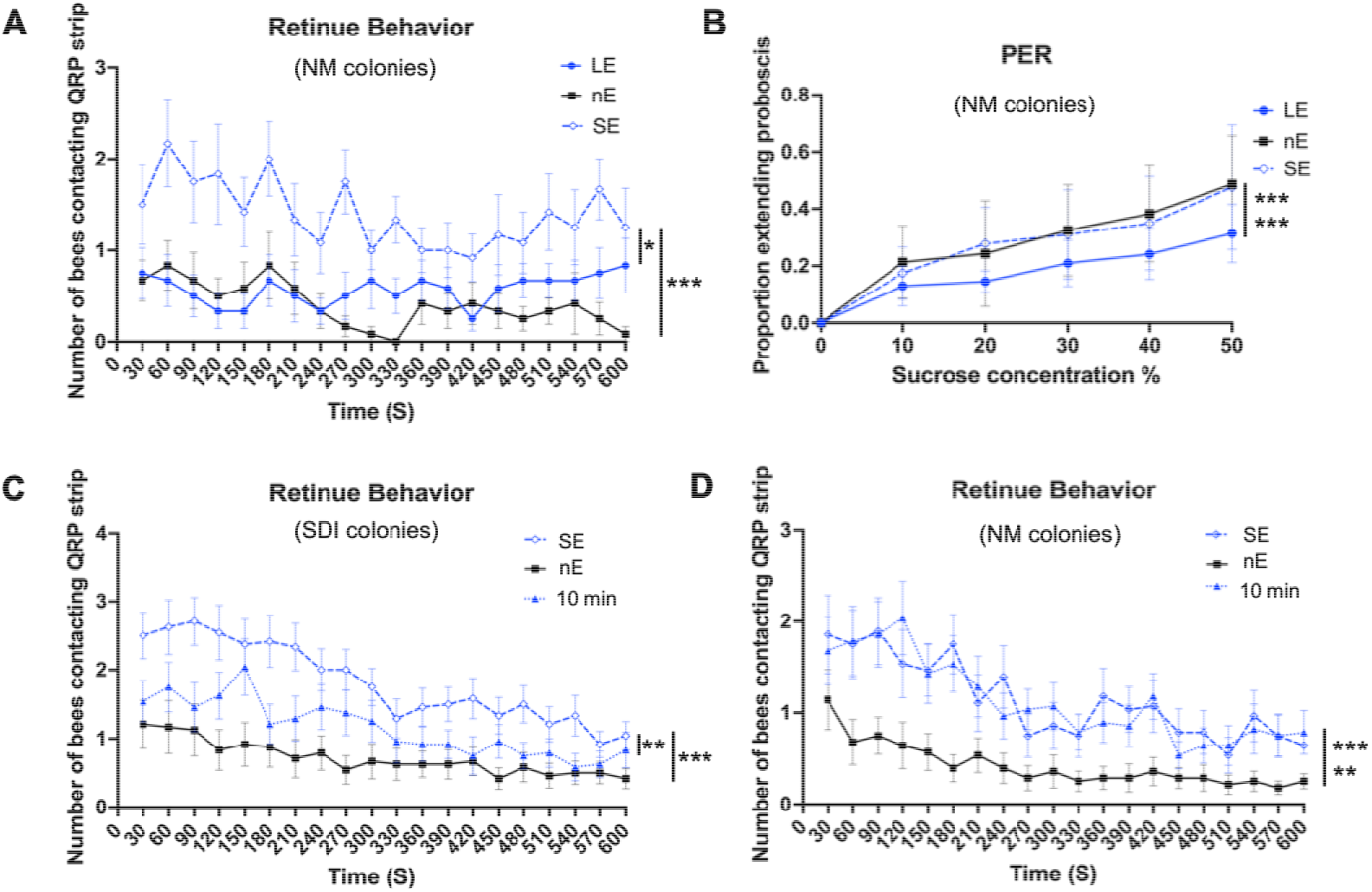
Effects of QRP stimulus strength on honey bee behavioral responses. **(A)** Retinue behavior of bees exposed to the weaker QRP stimulus (1h, SE) or the stronger QRP stimulus (3 days, LE) and of unexposed controls (nE). The number of bees contacting the QRP strip was recorded every 30 s during a 10-min assay. Data were collected in 2024 from 3 colonies, with 180 bees tested per treatment group. SE bees showed stronger retinue responses than LE (*p < 0.05) and nE bees (***p < 0.001). Individuals in these experiments were derived from naturally mated queens. **(B)** Proboscis extension response (PER) to increasing sucrose concentrations (0%–50%) in SE, LE, and nE bees, shown as the proportion of bees extending the proboscis. n = 127 (LE), 107 (nE), and 126 (SE) bees from 3 colonies (2024). LE bees were less responsive to sucrose than SE and nE bees (***p < 0.001 for each comparison). Individuals in these experiments were derived from naturally mated queens. (**C, D**) Retinue behavior of bees exposed to QRP for 1 h (SE), exposed for 10 min and then kept without QRP for 50 min (10 min), or not exposed to QRP (nE), measured as in (A). Data were collected in 2025, with 360 and 420 bees tested per treatment group from 6 SDI colonies (C) and from 6 NM colonies (D), respectively. In (C), bees in the 1-hour group showed stronger retinue responses than those in the 10-min group (**p< 0.01) and Control group (***p < 0.001). In (D), bees in both the 1-hour and 10-min groups showed stronger retinue responses than those in the Control group (***p < 0.001 and **p < 0.01, respectively). Bees in (C) were derived from queens each instrumentally inseminated with semen from a single drone, whereas bees in (D) were derived from naturally mated queens.

**Figure S2.**
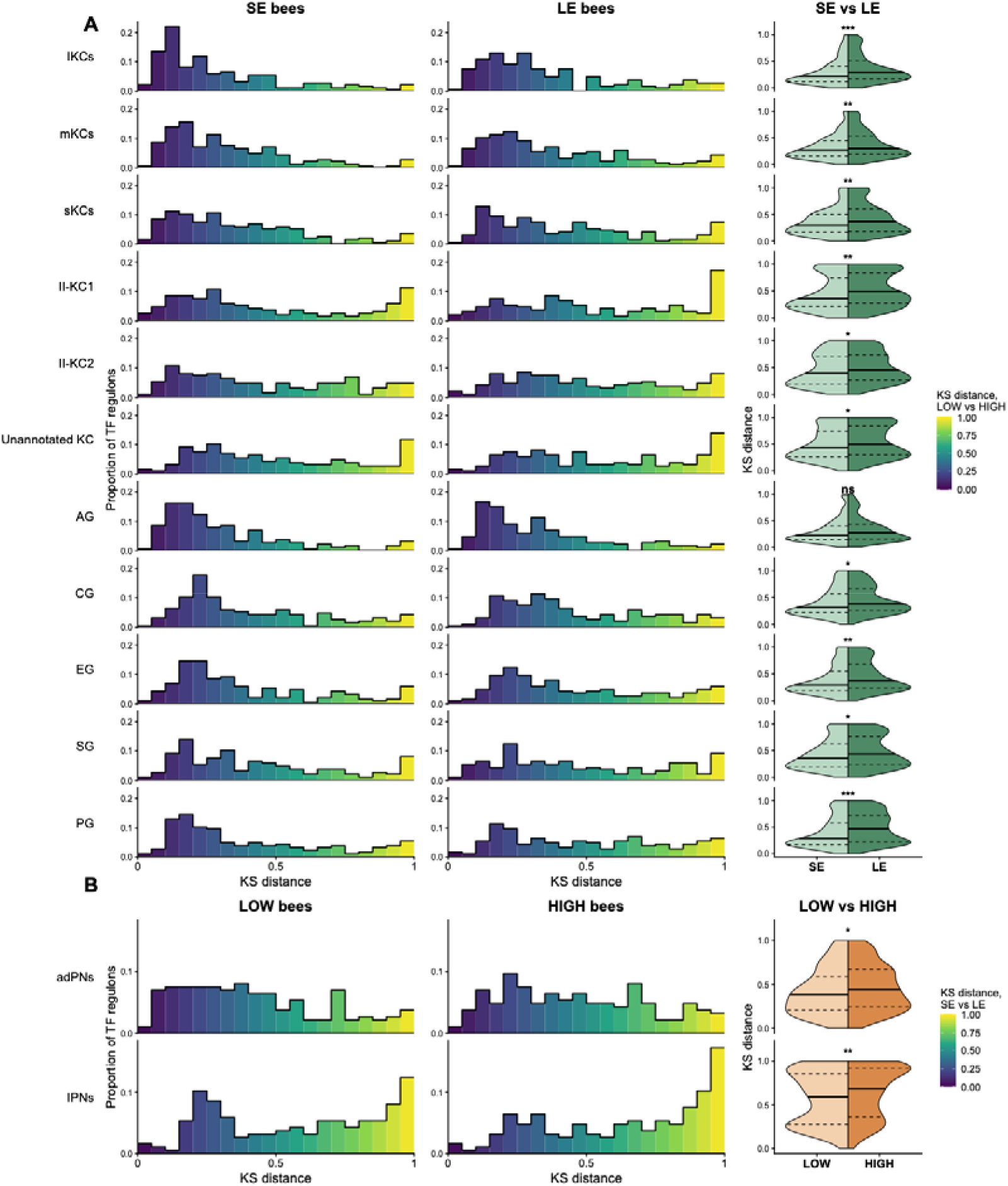
Expanded view of TF regulon regulatory dissimilarity across selected brain cell types shown in Figure 3. Regulatory dissimilarity was quantified for each TF regulon as the Kolmogorov–Smirnov (KS) distance between the distributions of SimiC regulon activity scores in two groups of bees. **(A)** KS distances between LOW- and HIGH-responsive bees, shown separately for SE and LE bees, across mushroom body Kenyon cell and glial subtypes. **(B)** KS distances between SE and LE bees, shown separately for LOW- and HIGH-responsive bees, across olfactory projection neuron subtypes. Left and middle columns: histograms of KS distances. Bar height indicates the proportion of TF regulons in each cell type whose KS distance falls within each bin, and bar color indicates KS distance. Right column: split violin plots comparing KS distance distributions between groups (A: SE vs LE; B: LOW vs HIGH). Solid lines indicate medians, and dashed lines indicate the 25th and 75th percentiles. Differences were tested using two-sided paired Wilcoxon signed-rank tests on TF regulons present in both groups (*p < 0.05, **p < 0.01, ***p < 0.001; ns, not significant). Abbreviations: lKCs, mKCs, and sKCs, large-, middle-, and small-type class I Kenyon cells; II-KC, class II Kenyon cells; AG, astrocytes glia; CG, cortex glia; EG, ensheathing glia; SG, subperineurial glia; PG, perineurial glia; adPNs, anterodorsal projection neurons; lPNs, lateral projection neurons; SE, weaker QRP stimulus (1-hour exposure); LE, stronger QRP stimulus (3-day exposure); LOW and HIGH, bees with low and high behavioral responsiveness to QRP, respectively.

**Figure S3.**
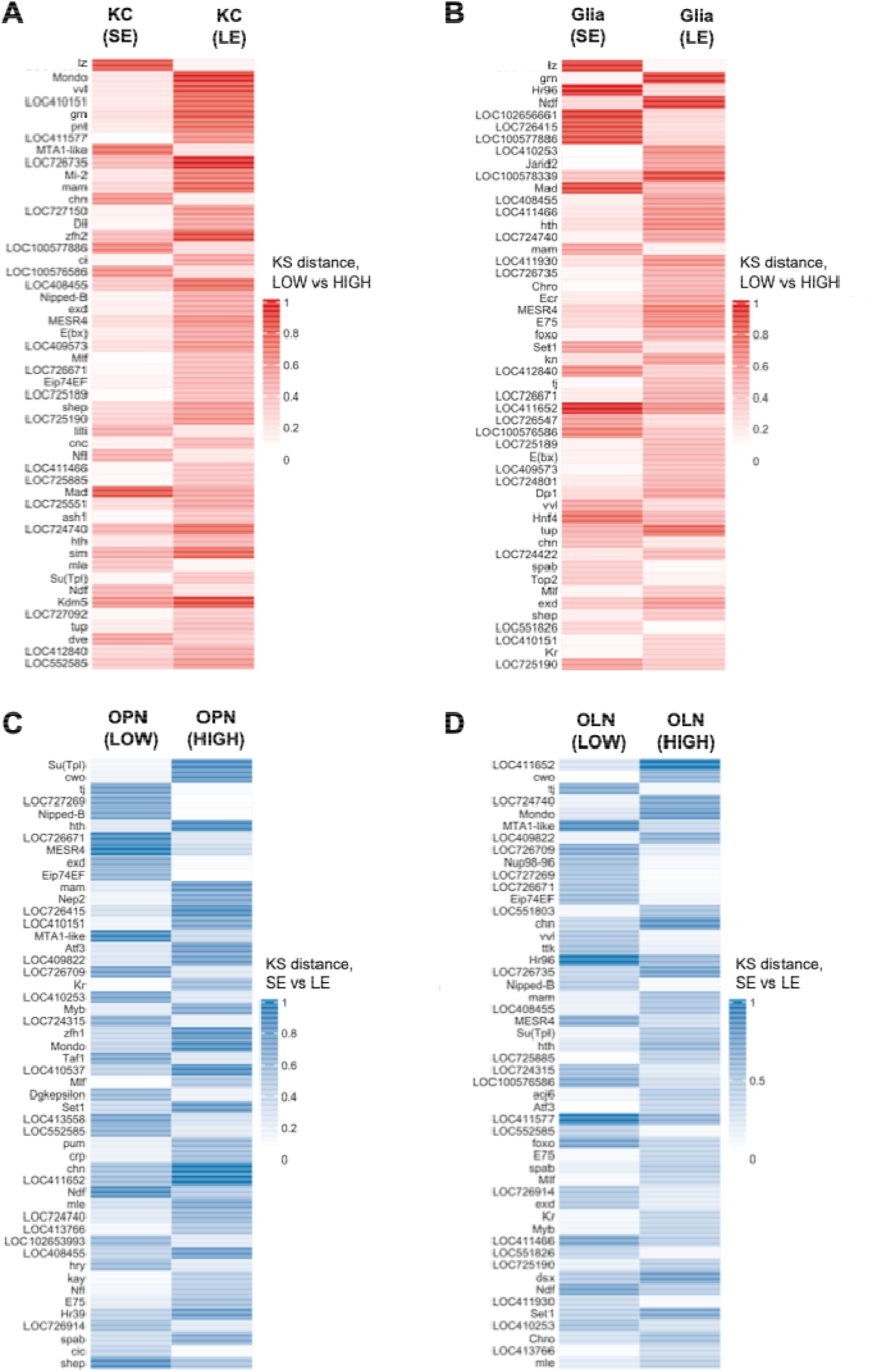
Transcription factors showing the largest differences in GRN regulatory dissimilarity across selected brain cell types. **(A and B)** Heatmaps showing the 50 transcription factors (TFs) with the largest differences in GRN regulatory dissimilarity between weaker and stronger QRP stimuli (SE and LE) in Kenyon cells (KCs; A) and glia (B). TFs were ranked by the absolute difference in Kolmogorov–Smirnov (KS) distance between the two exposure strengths, |KS(SE) − KS(LE)|. For each TF, the two columns show the Low-vs High-responsive bees (LOW and HIGH) KS distance after weaker and stronger QRP stimuli. Larger KS distances, indicated by increasing red intensity, represent greater regulatory dissimilarity between LOW and HIGH bees. **(C and D)** Heatmaps showing the 50 TFs with the largest differences in GRN regulatory dissimilarity between LOW and HIGH bees in olfactory projection neurons (OPNs; C) and optic lobe neurons (OLNs; D). TFs were ranked by the absolute difference in KS distance between the two behavioral-response groups, |KS(LOW) − KS(HIGH)|. For each TF, the two columns show the SE vs LE KS distance within LOW and HIGH bees. Larger KS distances, indicated by increasing blue intensity, represent greater regulatory dissimilarity between the two QRP stimulus strengths.

## Notes

### Competing Interest Statement

The authors have declared no competing interest.

### Funding

- Illinois Sociogenomics Initiative

